# OPTIMIS: Optimizing Personalized Therapies through Integrated Multiscale Intelligent Simulation

**DOI:** 10.64898/2026.03.24.713941

**Authors:** Zhaoqian Su, Yinghao Wu

## Abstract

Controlling complex biological systems across multiple scales remains a major challenge in computational medicine, because whole-body disease behavior is closely shaped by noisy cellular events at much smaller scales. Standard deterministic models often miss this molecular variability, while fully stochastic simulations are too slow for the repeated, high-throughput interactions needed to train artificial intelligence. To address this problem, we developed a new AI-based framework that combines a discrete stochastic Gillespie algorithm for microscale receptor dynamics with continuous, nonlinear ordinary differential equations for systemic macroscale behavior. To reach the speed needed for deep reinforcement learning (RL), we compress this hybrid system into a differentiable Neural ODE surrogate that acts as a fast digital twin. As a proof of concept, we applied this framework to engineered cellular therapy and used RL agents to learn dynamic, closed-loop treatment policies inside the surrogate environment. By tracking microscopic, unpredictable cellular activity as an early-warning signal, the AI learned to continuously adjust the drug dose—anticipating and stopping dangerous immune reactions before they could spiral out of control. This computational advance improved successful control rates to more than 70% in highly unstable simulated phenotypes and provides a practical, general framework for adaptive intervention in multiscale biological systems.

## Introduction

CAR-T cell therapy is a powerful immunotherapy in which engineered T cells are used to recognize and eliminate malignant cells [1]. Despite its clinical success, treatment response is highly variable across patients, and outcomes are shaped by interacting processes across multiple biological scales [2]. At the cellular level, receptor binding and synapse activation regulate how effectively CAR-T cells engage tumor cells [3]. At the tissue and systemic levels, these interactions influence tumor killing, CAR-T expansion or exhaustion, and inflammatory cytokine release [4]. One of the major clinical challenges is balancing efficacy against toxicity, especially cytokine release syndrome–like behavior, where excessive immune activation can produce dangerous systemic inflammation [5]. Drug modulation adds another layer of complexity. Agents such as Dasatinib can alter signaling and receptor activity, potentially providing a mechanism to tune CAR-T activity dynamically [6]. In principle, this could improve safety by dampening toxic over-activation, or improve efficacy by shaping receptor-state dynamics over time [7]. In practice, however, the treatment design problem is difficult because the relevant biology is nonlinear, patient-dependent, and strongly multiscale [8]. Static dosing schedules are unlikely to be optimal across heterogeneous patients or across different stages of treatment response.

This motivates the need for computational platforms that can integrate mechanistic biology with adaptive decision-making. A useful platform must represent both microscale biology, such as receptor activation state, and macroscale disease trajectories, such as tumor burden and cytokine dynamics. It must also support sequential treatment optimization under uncertainty, where the best decision at one time point depends on the evolving state of the patient and the downstream consequences of current dosing. Existing computational approaches, such as Quantitative Systems Pharmacology (QSP) [9–17] and standard pharmacokinetic/pharmacodynamic (PK/PD) models [18–24], have provided valuable baseline insights into CAR-T population dynamics and tumor clearance. However, these frameworks predominantly rely on deterministic, macroscale differential equations that fail to capture the highly volatile, stochastic molecular jitter of receptor-antigen binding at the immunological synapse [25]. Conversely, strictly discrete stochastic frameworks [26–34] that do capture this cellular-level noise are computationally intractable when scaled to simulate systemic patient trajectories over a multi-week clinical horizon. Furthermore, treatment optimization within these traditional models is typically limited to evaluating pre-defined, static dosing schedules, rather than discovering proactive, closed-loop control strategies [35, 36]. Consequently, there remains a critical computational bottleneck: translating the rigorous, multiscale biophysics of the CAR-T synapse into an environment fast enough, and scalable enough, to train advanced adaptive decision-making algorithms.

To overcome these computational bottlenecks, we introduce a novel, AI-driven framework OPTIMIS. OPTIMIS utilizes a slow-fast coupled hybrid architecture that rigorously integrates the stochastic molecular jitter of the immunological synapse—modeled via a discrete Gillespie algorithm—with the continuous, deterministic differential equations governing systemic tumor clearance and cytokine release. To achieve the execution speeds required for reinforcement learning, we distill this complex hybrid system into a differentiable Neural Ordinary Differential Equation (NeuroODE) surrogate that approximates macro-scale dynamics while remaining coupled to mechanistic receptor-state updates. This surrogate environment allows us to deploy deep Reinforcement Learning (RL) agents capable of discovering dynamic, closed-loop pharmacological interventions that adapt to a patient’s evolving biophysical state in real-time.

Our preliminary results demonstrate that this multiscale, RL-driven approach fundamentally outperforms standard clinical heuristics, particularly for high-risk patient phenotypes. By training the AI on a longitudinal synthetic cohort of 240 patients, the agent autonomously discovered a proactive, 3-phase “surfing” policy for aggressive disease profiles: a preemptive Dasatinib brake to suppress initial hyper-activation, a controlled taper to allow steady tumor clearance, and a highly reactive “soft landing” pulse to prevent toxic cascades. Where standard reactive clinical protocols resulted in 0% success and universally lethal cytokine storms in high-risk simulated patients, our multiscale RL agent achieved up to a 74% curative success rate. By leveraging microscale receptor dynamics as an early-warning computational biomarker, the AI successfully eradicated the tumor burden while mathematically pinning systemic cytokines strictly below life-threatening safety thresholds.

## Model and Methods

### Summary of computational framework

We developed a multiscale simulation-and-control framework for adaptive CAR-T therapy optimization under drug modulation, called OPTIMIS (Optimizing Personalized Therapies through Integrated Multiscale Intelligent Simulation), as illustrated in **Figure 1**. The framework couples a mechanistic microscale synapse model, a learned macro-scale neural ODE surrogate, and a reinforcement learning controller inside a persistent training pipeline. At the microscale, receptor activation dynamics are modeled by simulating synapse-state transitions as a function of baseline binding kinetics, inactive and active receptor counts, drug exposure, and cytokine level. This component provides a mechanistic update of receptor activity at each simulated treatment step. At the macro-scale, tumor burden, CAR-T cell abundance, and cytokine concentration are represented by a drug-aware neural ODE. To train this surrogate, we first generated a synthetic cohort dataset in which each simulated patient was assigned to a standard or aggressive phenotype with different baseline receptor-binding parameters. For each patient, a stochastic dosing schedule was sampled. The resulting longitudinal dataset contained tumor burden, CAR-T level, cytokines, receptor activation, drug exposure, and phenotype labels. A neural ODE was then trained on these transitions so that it could serve as a differentiable, computationally efficient digital twin of the macro-scale system.

**Figure 1:**
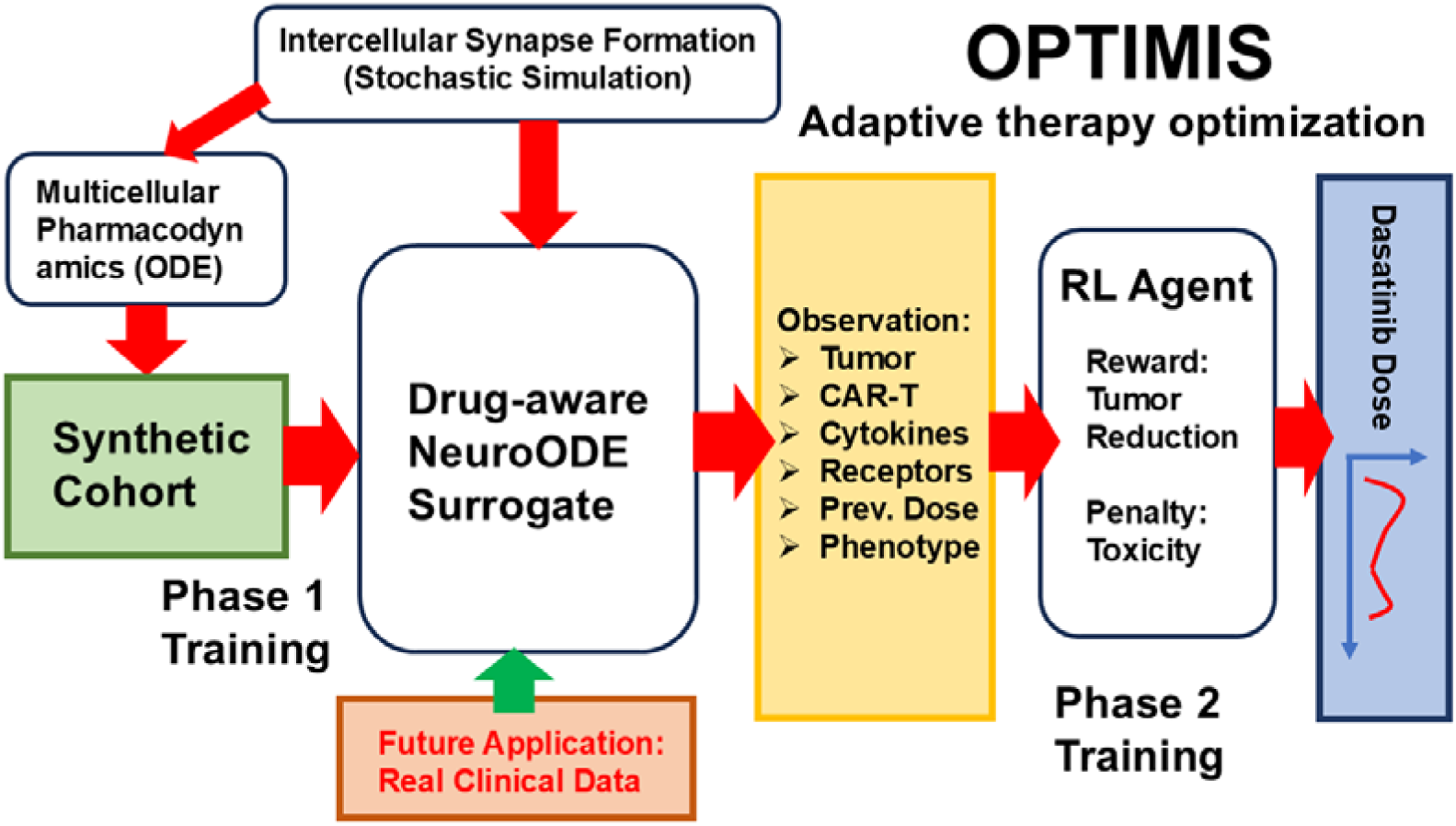
The computational pipeline is divided into two primary phases. A mechanistic multiscale model, integrating stochastic intercellular synapse simulations with multicellular pharmacodynamics, generates a synthetic patient cohort. This longitudinal data is used to train a computationally efficient, drug-aware Neural ODE surrogate, which serves as a macro-scale digital twin. A reinforcement learning (RL) agent interacts with the trained Neural ODE environment. At each decision step, the agent receives a multi-dimensional state observation (including tumor burden, CAR-T cell count, cytokine levels, and phenotype hints) and outputs an optimized, continuous Dasatinib dose. The RL reward function is structured to maximize tumor reduction while explicitly penalizing systemic toxicity and abrupt dosing changes. The architecture also highlights a translational pathway for future integration of real-world clinical data to iteratively refine the surrogate model.

The treatment optimization layer was formulated as a reinforcement learning problem. The RL environment used the trained neural ODE to create a multiscale digital twin. At each decision step, the agent observed a compact state vector composed of normalized tumor burden, CAR-T count, cytokines, receptor activation, recent temporal changes in tumor and cytokines, previous dose, treatment-time fraction, and a phenotype hint. The action was a continuous Dasatinib dose in the range from zero to one. Given an action, receptor activation was first updated through the mechanistic synapse simulator, and the macro-state was then advanced with the neural ODE. The reward function was designed to promote tumor reduction while penalizing systemic toxicity, cytokine escalation, excessive drug exposure, and abrupt changes in dose.

In summary, the developed method is a multiscale adaptive control framework in which a mechanistic receptor-level simulator informs a learned system-level digital twin, and that digital twin is used as the environment for reinforcement learning of personalized, phenotype-aware dosing strategies.

### Multiscale Hybrid Modeling of CAR-T Pharmacodynamics

As illustrated in **Figure 2**, a computationally efficient hybrid model was first built to model CAR-T pharmacodynamics. This architecture isolates the rapid, stochastic noise of the immunological synapse at the cellular microscale, utilizing its output to drive the continuous, deterministic differential equations governing systemic macroscale disease progression. The systemic biological state is modeled via a system of coupled, non-linear ordinary differential equations (ODEs). Let T represent the tumor cell burden, C the circulating CAR-T cell population, and I the systemic inflammatory cytokine concentration. The dynamics are fundamentally driven by α (alpha), the dynamic fraction of actively engaged CAR receptors computed by the microscale model. We define a set of intrinsic biological rate parameters to govern these interactions.

**Figure 2:**
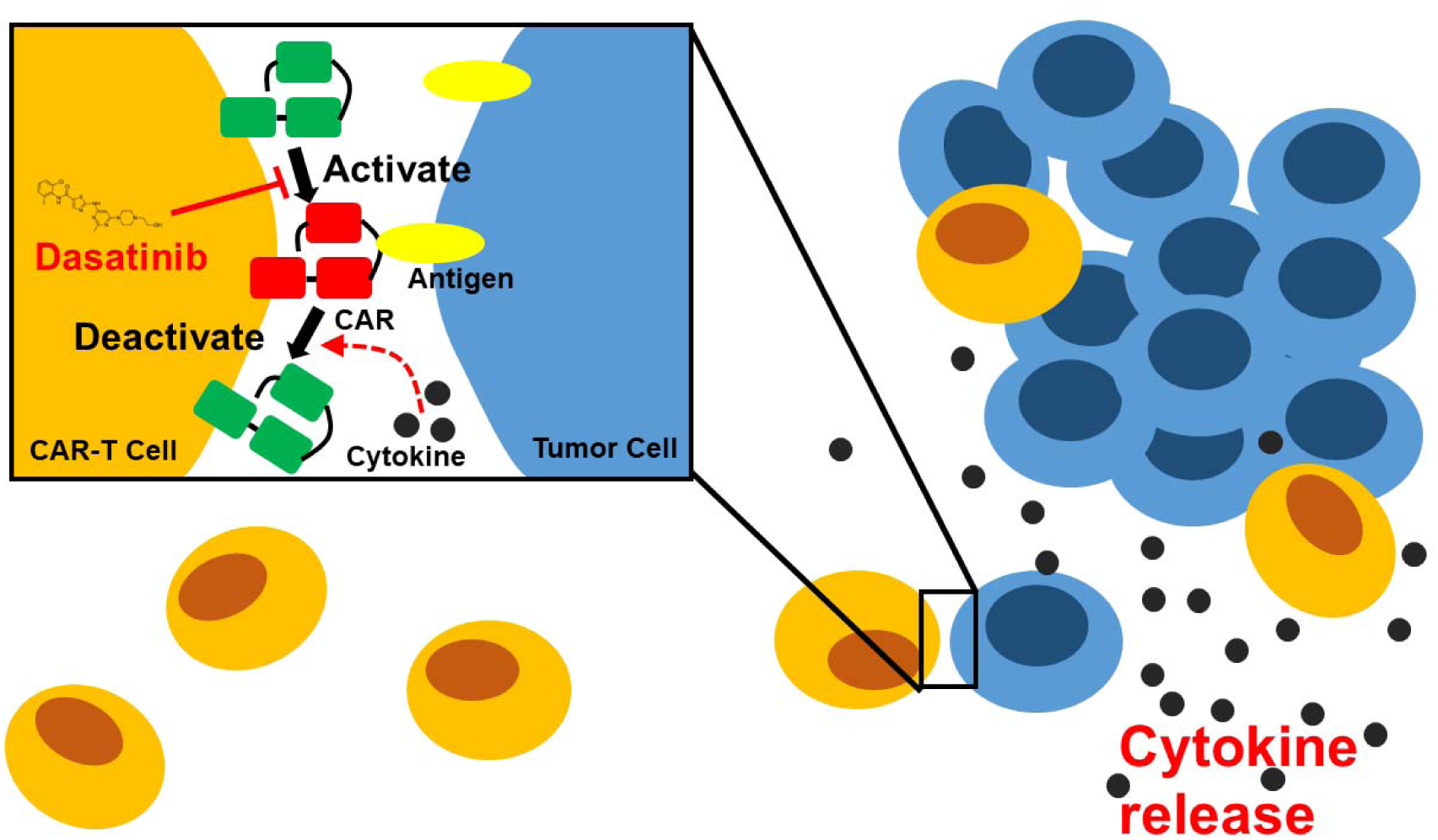
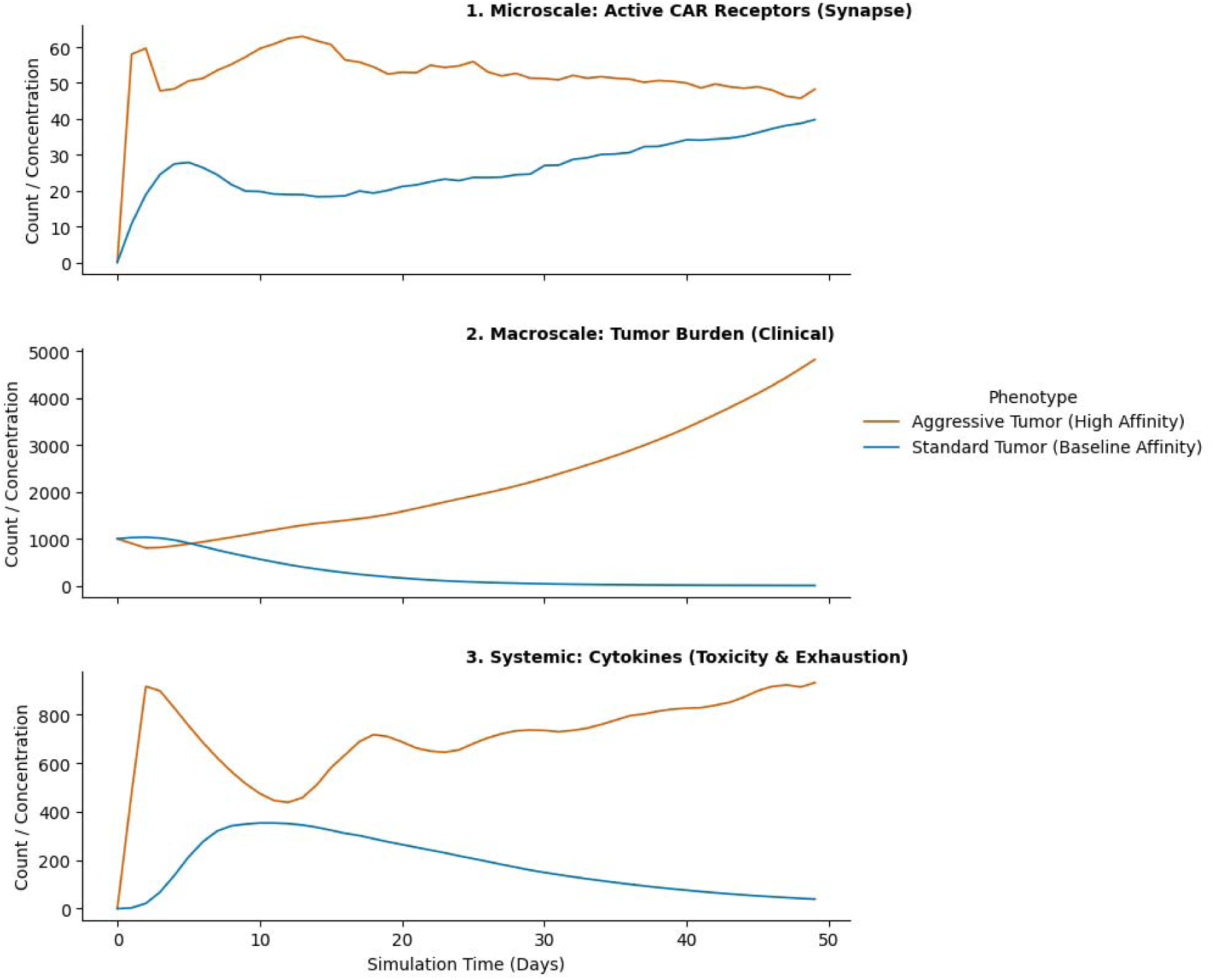
The multiscale dynamics of the CAR-T immunological synapse and Dasatinib intervention. At the cell-cell interface (small inset), tumor surface antigens engage with Chimeric Antigen Receptors (CARs), driving receptor activation to initiate the cytotoxic T-cell response. To mitigate the risks of hyper-activation, the tyrosine kinase inhibitor Dasatinib acts as a dynamic pharmacological “off-switch” by directly blocking this critical activation step. If left uninhibited, sustained receptor engagement not only drives targeted tumor cell clearance but also triggers the prolific release of inflammatory cytokines; these cytokines subsequently fuel a regulatory feedback loop that accelerates CAR deactivation and ultimately contributes to systemic immune exhaustion.

Tumor Kinetics: Tumor progression is governed by a baseline exponential growth rate (r_T) and active clearance mediated by engaged CAR-T cells at a targeted killing rate (k_k):

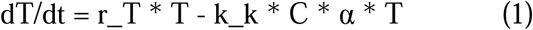

CAR-T Cell Expansion and Exhaustion: The CAR-T population expands at a proliferation rate (p_C) in response to target engagement but undergoes natural clearance (d_C). Crucially, we mathematically model exhaustion and activation-induced cell death via a non-linear storm penalty (S_p):

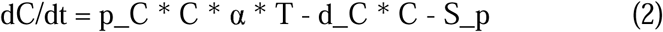

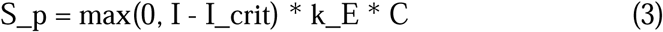

This penalization ensures that if systemic inflammation crosses a critical, life-threatening threshold (I_crit), the CAR-T cells rapidly lose viability scaled by an exhaustion rate (k_E), enforcing a mathematical limit on hyper-activation.

Systemic Cytokine Production: Cytokine release is modeled as a highly non-linear amplification cascade. Production scales with the cube of the active receptor fraction at a distinct rate (p_I), mirroring the exponential biological reality of immune signaling, alongside a natural systemic clearance rate (d_I):

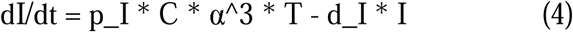

At the microscale, the immunological synapse is modeled as a continuous-time Markov chain using the Gillespie Stochastic Simulation Algorithm (SSA). This captures the discrete, noisy toggling of individual CAR receptors between inactive (R_inact) and active (R_act) states. The baseline transition propensities are dynamically coupled to the real-time macroscale environment, specifically the administered Dasatinib drug dose (D, ranging from 0 to 1) and systemic cytokines (I):

Activation Rate (k_bind): The binding propensity is suppressed by the targeted administration of the tyrosine kinase inhibitor:

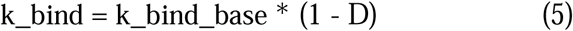

Deactivation Rate (k_unbind): Receptor shedding and synapse destabilization are positively correlated with a hostile, highly inflammatory systemic environment [37], driven by a cytokine feedback scaling parameter (γ):

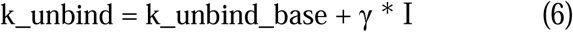

By determining the exact time to the next molecular event via an exponential distribution based on the total system propensity, the SSA accurately simulates the inherent molecular jitter of the off-switch mechanism.

To prevent mathematical stiffness and ensure rapid execution during AI training, the model employs a strict time-scale separation (“handshake”) protocol. At the beginning of each macroscopic integration step (dt), the ODE solver halts and passes the current I and D values to the Gillespie SSA. The microscale model rapidly simulates the requested time window and returns the absolute integer count of active receptors. This value is normalized to derive α, which is then assumed constant by the ODE solver for the duration of that specific macro-step. This discrete-continuous handshake maintains rigorous biophysical fidelity while achieving the millisecond execution speeds required for high-throughput reinforcement learning.

### Synthetic data generation

A virtual cohort is simulated over a fixed treatment horizon. Each patient is assigned to either a standard or aggressive phenotype. The phenotype determines baseline receptor-binding properties, which in turn affect the outputs of the synapse model. Initial tumor burden, CAR-T count, cytokine level, and receptor activity are sampled from distributions to reflect inter-patient variability. For each patient, a drug schedule is generated stochastically from a family of patterns that includes low-dose constant regimens, moderate constant regimens, pulsatile schedules, upward ramps, downward ramps, and random-walk-like profiles. This ensures that the training data expose the system to a broad range of treatment trajectories.

At each simulated day, the current drug dose is first passed into the microscale synapse model along with the patient’s current cytokine level and receptor counts. The synapse model returns the updated receptor activation state. That receptor state is then passed into a macro-scale ordinary differential equation system governing tumor burden, CAR-T abundance, and cytokines. In this way, the synthetic data are generated by alternating between microscale signaling updates and macro-scale population-level evolution. The resulting dataset records, for each patient and time point, tumor burden, CAR-T counts, cytokine concentration, receptor activation, drug exposure, phenotype label, and baseline kinetic parameters.

This stage is critical because it creates consistent multiscale trajectories in which drug exposure affects receptor activation, receptor activation affects macro-scale tumor-immune dynamics, and phenotype modulates the entire response profile. The generated dataset therefore functions as a structured set of state transitions from which the macro-scale surrogate can be learned. In practical terms, this stage also establishes the digital twin’s training distribution, so the quality and diversity of these synthetic trajectories strongly influence the robustness of the downstream controller.

### Learned macro-scale neural ODE surrogate

A neural ordinary differential equation [38–40] is used as a learned macro-scale surrogate. This module serves as a differentiable and computationally efficient digital twin of the system-level disease dynamics. Its role is to replace repeated direct simulation of hand-crafted macro-scale equations during control training, while remaining coupled to microscale receptor-state information.

The neural ODE takes as state input tumor burden, CAR-T cell abundance, cytokine concentration, normalized receptor activation, and drug level. Internally, the model is implemented as a multilayer neural network that predicts the time derivative of the macro-scale variables. Receptor fraction and drug level are treated as control-context variables with zero dynamics in the surrogate, while the network learns the macro-scale derivatives for tumor, CAR-T population, and cytokines. This structure makes the model “drug-aware” and “receptor-aware” without forcing it to predict the receptor kinetics directly, since those are already handled by the microscale synapse simulator.

Training is performed on the synthetic multiscale transition dataset. Pairs of consecutive states are extracted from each patient trajectory, with receptor activation normalized to a fraction. The neural ODE is then optimized so that integrating the learned dynamics forward over one time unit reproduces the next observed macro-state. SmoothL1 loss is used to fit the predicted next-step tumor, CAR-T, and cytokine values. Because the model learns continuous-time dynamics rather than a purely discrete mapping, it is well suited for integration into downstream control settings where sequential updates and differentiability are valuable.

In the final framework, the neural ODE acts as the macro-scale propagation engine inside the RL environment. After the synapse model updates receptor activation, the neural ODE advances tumor burden, CAR-T count, and cytokines to the next time step. This makes the digital twin computationally lighter than a fully mechanistic simulator at all scales, while retaining a biologically meaningful coupling between receptor regulation and systemic response. The neural ODE therefore provides the balance between flexibility, speed, and multiscale structure that makes repeated control optimization feasible.

### Reinforcement learning controller for adaptive dosing

The last step of the framework is the reinforcement learning controller, which transforms the multiscale digital twin into a sequential decision-making environment for adaptive therapy optimization. This component is responsible for learning how to choose drug doses over time in order to maximize therapeutic benefit while minimizing toxicity and unnecessary treatment exposure.

The environment state is constructed from normalized system variables and recent temporal context. The observation includes tumor burden, CAR-T count, cytokine level, receptor activation, recent change in tumor burden, recent change in cytokine level, previous dose, fraction of the treatment horizon elapsed, and a phenotype hint indicating standard or aggressive disease. The action is a continuous scalar dose between zero and one, representing the amount of drug to administer at the current step. To encourage smoother policies, the effective dose applied to the system is filtered using the previous action so that treatment changes are gradual rather than impulsive.

Each environment step proceeds in two stages. First, the chosen dose is passed to the microscale synapse simulator, which updates receptor activation as a function of the current state. Second, the updated receptor state and drug level are passed to the neural ODE surrogate, which advances the macro-scale disease variables by one time unit. This produces the next simulated patient state. In this way, the RL controller does not interact with a single black-box transition function; it interacts with a structured multiscale digital twin in which drug action is routed through receptor biology and then through system-level disease evolution.

The reward function is designed to encode the clinical tradeoff between efficacy and safety. The agent is rewarded for reducing tumor burden and making daily progress in tumor shrinkage. It is penalized for large residual tumor burden, elevated cytokine toxicity above a soft threshold, rapid cytokine increases, total drug usage, and abrupt changes in dose. Episodes terminate early if cytokine levels exceed a dangerous threshold, corresponding to a toxicity crisis. Episodes also terminate successfully when tumor burden is driven below a low threshold while cytokines remain in a safe range. If neither event occurs, the episode ends at the maximum treatment horizon with an additional penalty for residual disease.

The controller is trained using proximal policy optimization (PPO) [41, 42]. A neural feature extractor processes the observation vector before policy and value heads produce the action distribution and value estimate. PPO is well suited to this task because it supports stable training in continuous action spaces and handles noisy, nonlinear environments effectively.

Overall, this RL component converts the multiscale simulation framework into an adaptive treatment optimization platform. Instead of prescribing a fixed dosing schedule, it learns a closed-loop control law that conditions each decision on the evolving tumor, immune, cytokine, receptor, and phenotype state of the simulated patient.

## Results

### Generation of patient cohort

To train and validate the Reinforcement Learning (RL) agent, we first generated a highly detailed, 12,000-point longitudinal synthetic dataset using our hybrid mechanistic model. The data comprised an evenly balanced cohort of 240 simulated patients tracked over a 50-day clinical horizon. Patients were categorized into two distinct phenotypes: a “Standard” cohort (N=120, baseline binding rate k_bind_base≈0.1) and an “Aggressive” cohort (N=120, k_bind_base≈0.9).

As visualized in the representative trajectory (**Figure 3**), the Standard cohort (blue curves) typically achieved steady tumor clearance with manageable cytokine release. Conversely, the Aggressive cohort (orange curves) exhibited hyper-proliferation; while tumor clearance was rapid, unmanaged CAR-T expansion (peaking at ∼978 cells/uL) consistently triggered lethal Grade 3/4 cytokine storms. In the worst unmanaged cases, systemic cytokines spiked to 1,687 pg/mL—more than three times our defined lethal toxicity threshold of 500 pg/mL, necessitating precise, dynamic pharmacological intervention.

**Figure 3:**
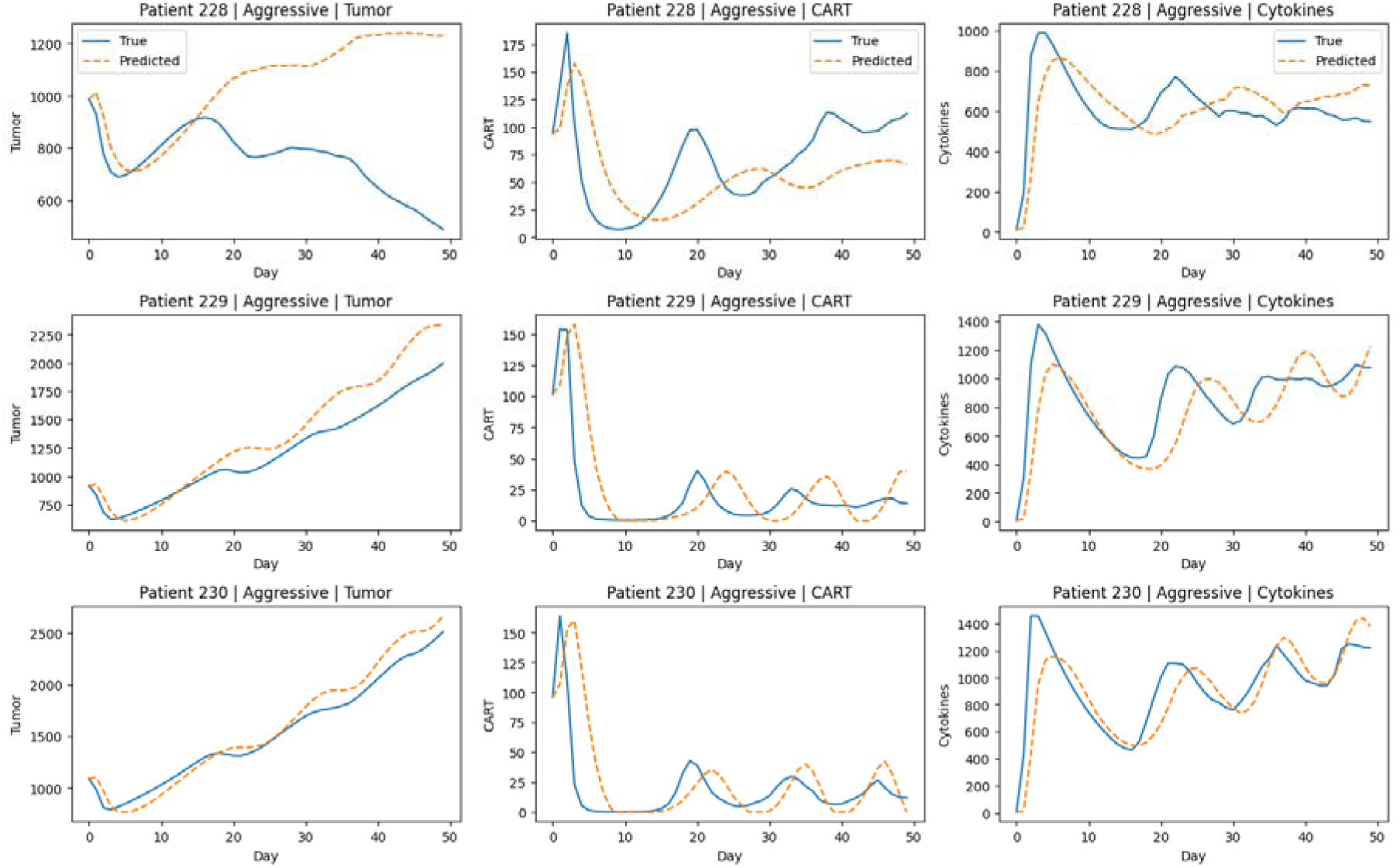
The generation of synthetic patient trajectories. The simulation produces multiscale time-series data across distinct simulated phenotypes: a Standard Tumor (blue) and an Aggressive Tumor (orange). (Upper panel) Microscale: Stochastic fluctuations in active CAR receptors at the immunological synapse. (Middle panel) Macroscale: Clinical tumor burden progression over a 50-day synthetic trajectory. (Lower panel) Systemic: Inflammatory cytokine concentrations. While the synthetic standard phenotype resolves without intervention, the aggressive phenotype generates a complex failure trajectory characterized by rapid receptor hyper-activation, a severe cytokine storm, and subsequent CAR-T exhaustion.

### High-fidelity surrogate modeling via NeuralODEs

Because the discrete stochastic Gillespie algorithm is computationally expensive, we trained a deep Neural Ordinary Differential Equation (NeuroODE) to act as a differentiable digital twin. The surrogate takes as input tumor burden, CAR-T abundance, cytokine concentration, normalized receptor activation, and drug exposure, and predicts the continuous-time evolution of the macro-scale system over each treatment interval. In the OPTIMIS framework, receptor activation remains mechanistically updated by the microscale synapse simulator, while the NeuroODE propagates the larger-scale disease state forward in time. This design preserves biologically meaningful coupling while greatly reducing computational cost during repeated policy optimization.

As shown in the validation curves from three randomly selected patients (**Figure 4**), the NeuroODE learned the local transition structure of the simulated environment with high accuracy. Across 11,760 one-step transitions, the surrogate achieved low normalized mean absolute error (NMAE) for all three macro variables: 0.0017 for tumor burden, 0.0048 for CAR-T abundance, and 0.0105 for cytokines. The corresponding one-step root mean squared errors were 15.26 for tumor, 9.84 for CAR-T, and 56.94 for cytokines. These results indicate that the surrogate accurately captures the short-horizon state updates most relevant to sequential control.

**Figure 4:**
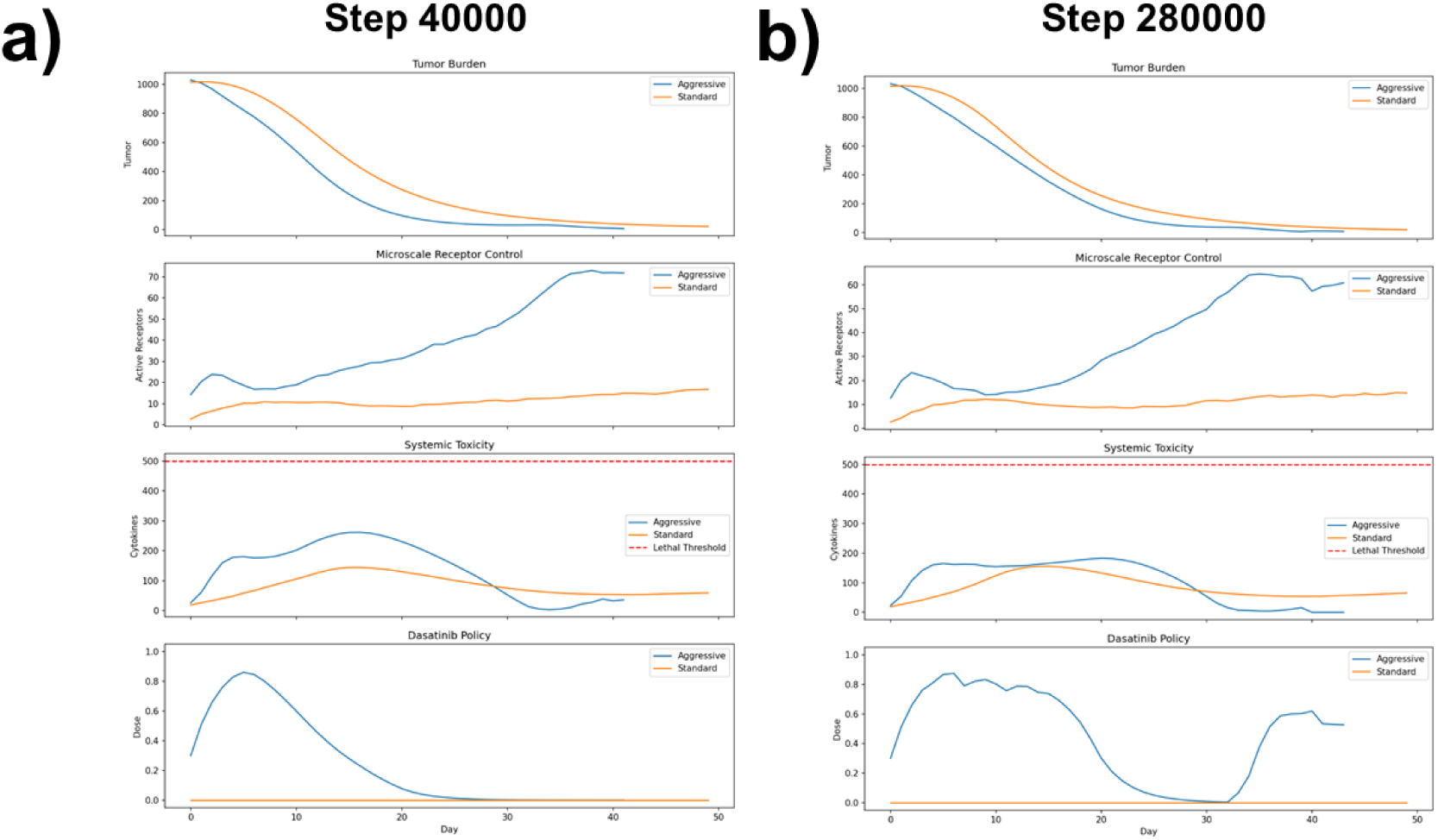
The hybrid Gillespie-ODE architecture was distilled into a predictive surrogate model. This figure demonstrates the surrogate’s predictive accuracy across three distinct simulated patients presenting with the highly volatile Aggressive tumor phenotype. Across all three cohorts, the predicted trajectories (dashed orange) for macroscopic Tumor Burden, CAR-T population dynamics, and systemic Cytokine concentrations closely track the ground-truth data (solid blue) over a 50-day period. The surrogate is able to captures the complex, non-linear oscillatory behavior of the CAR-T expansion and cytokine release cycles, ensuring the RL agent is trained in a high-fidelity environment that accurately mirrors the underlying biological mechanics.

To further assess fidelity beyond one-step prediction, we evaluated free multi-step rollouts on 12 held-out aggressive patients. In this setting, the model preserved the qualitative structure of the trajectories, including early tumor decline, CAR-T expansion and contraction, and the appearance of inflammatory cytokine surges. However, the rollout analysis also revealed progressive long-horizon drift. Mean rollout MAE across held-out patients was 243.67 for tumor, 25.44 for CAR-T, and 155.79 for cytokines, corresponding to rollout NMAE values of 0.0408, 0.0260, and 0.0924, respectively. Thus, while the NeuroODE reproduced the broad trajectory morphology of the multiscale system, it was less accurate as a long-term patient-specific forecaster than as a short-step transition model.

This distinction is important for interpreting the role of the surrogate within OPTIMIS. The current NeuroODE behaves most strongly as a locally accurate, control-oriented surrogate rather than as a fully faithful long-horizon digital twin of individual patients. For reinforcement learning, this level of fidelity is sufficient: the controller only requires a transition model that preserves short-term biological trends, treatment-response tradeoffs, and reward-relevant dynamics. Indeed, the low one-step errors indicate that the surrogate provides a stable and informative environment for policy learning. At the same time, the observed rollout drift highlights an important area for future refinement, including multi-step rollout training, richer patient conditioning, and eventual recalibration using experimental or clinical data. Overall, the NeuroODE achieves the key methodological objective of enabling fast, differentiable, multiscale control experiments while retaining biologically structured dependence on receptor activation and drug exposure. Its performance supports the use of OPTIMIS as a proof-of-concept computational platform for adaptive therapy design, while also clarifying that further improvements will be needed before claiming high-fidelity long-term patient forecasting.

### Evolution of the RL policy

The RL agent’s learning trajectory (**Figure 5**) demonstrated a profound shift from reactive survival to proactive, personalized medicine. Early Training (**Figure 5a**): Initial models exhibited a naive “survival” heuristic. To avoid the massive mathematical penalties associated with cytokine toxicity, the agent applied maximum Dasatinib braking continuously. While this prevented cytokine storms, it completely suppressed CAR-T activity, allowing the tumor burden to grow unchecked (reaching upwards of 5,969 cells), resulting in treatment failure. By step 280,000, the agent mastered the environment (**Figure 5b**). For the Standard cohort, the agent recognized the low risk and flatlined the drug dose at 0.0, allowing natural clearance. For the Aggressive cohort, the agent learned to execute a sophisticated, 3-phase “surfing” policy: a preemptive heavy brake (Dose ≈ 0.85) to suppress initial hyper-activation, a controlled taper to zero to allow tumor clearance, and a highly reactive “soft landing” pulse (Dose ≈ 0.6) at Day 35 to mathematically pin systemic cytokines just below the critical penalty warning threshold of 200 pg/mL.

**Figure 5:** This figure illustrates the RL agent’s learned Dasatinib administration strategy at an early training phase versus a mature, highly optimized phase for both Standard (orange) and Aggressive (blue) tumor phenotypes. For the Standard phenotype, the agent correctly learns to withhold the drug entirely (Dose = 0), as natural tumor clearance occurs without systemic cytokines crossing the lethal threshold (red dashed line). For the Aggressive phenotype, early training **(a)** yields a broad, generalized dosing curve that successfully prevents lethal toxicity but lacks precision. By late-stage training **(b)**, the agent discovers a highly dynamic, multi-phase intervention strategy. It applies a strong preemptive dose to blunt the initial microscale receptor hyper-activation and subsequent cytokine storm, strategically tapers the drug to permit CAR-T expansion and tumor clearance, and executes a secondary micro-dosing phase (Days 30–40) to suppress a late-stage inflammatory rebound.

These results suggest that the RL controller changed substantially over training. Early in training, the agent mainly learned a simple safety-driven behavior: it gave high Dasatinib doses to suppress CAR-T activity and avoid cytokine penalties. This reduced toxicity risk, but it also weakened treatment and often allowed tumor burden to remain high. In this stage, the policy behaved more like a general brake than a useful therapeutic strategy. Later in training, the controller became more selective and phenotype-aware. For standard patients, it learned that little or no drug was needed under the current model, and it therefore kept dosing near zero. For aggressive patients, however, it learned a more structured and adaptive pattern. The mature policy typically applied an early strong dose to prevent excessive initial activation, then reduced dosing to allow tumor killing, and in many cases added a later dose pulse to prevent delayed cytokine escalation. This produced a clear multi-phase strategy rather than a fixed schedule.

Importantly, this behavior was not manually programmed. It emerged from reinforcement learning as the agent optimized tumor control, toxicity, cumulative drug use, and dose smoothness at the same time. The final policy therefore behaved like a closed-loop control strategy: it adjusted treatment according to the evolving state of the simulated patient instead of following a single preset regimen. Overall, the training process showed a transition from blunt suppression to a more balanced and adaptive treatment policy, especially for the aggressive cohort.

### Ablation analysis: the necessity of multiscale and phenotypic features

We next performed an ablation analysis to determine which parts of the observation space were most important for successful control. Five model variants were evaluated, as listed in **Table 1**. Full Model used the complete observation vector, including tumor burden, CAR-T count, cytokines, receptor activation, short-term change features, previous dose, treatment-time fraction, and phenotype hint. No_Class_Hint removed the phenotype indicator that distinguishes standard from aggressive disease. No_Receptor_Channel removed the receptor-activation input from the observation. No_Delta_Features removed the short-term change variables for tumor burden and cytokines. No_Prev_Dose removed the previous-dose input, preventing the controller from explicitly tracking its recent intervention history.

**Table 1:** Ablation analysis demonstrating the critical contribution of individual neural network state features to the overall success rate and dosing efficacy of the reinforcement learning agent.

| *Policy* | *Cohort* | *Num_Patients* | *Success_Rate* | *Mean_Final_Tumor* | *Mean_Peak_Cytokines* | *Frac_Cytokines_Over_200* | *Frac_Cytokines_Over_300* | *Mean_Cumulative_Dose* |
| --- | --- | --- | --- | --- | --- | --- | --- | --- |
| *Full_Model* | Standard | 103 | 0 | 20.854806 | 165.751112 | 0.126214 | 0 | 0 |
| *Full_Model* | Aggressive | 97 | 0.742268 | 7.337489 | 195.653031 | 0.257732 | 0 | 15.882653 |
| *No_Class_Hint* | Standard | 103 | 0 | 20.92693 | 168.104941 | 0.165049 | 0 | 0 |
| *No_Class_Hint* | Aggressive | 97 | 0 | 231.108949 | 471.436828 | 1 | 1 | 1.896227 |
| *No_Delta_Features* | Standard | 103 | 0 | 21.136725 | 163.809805 | 0.07767 | 0 | 0 |
| *No_Delta_Features* | Aggressive | 97 | 0.773196 | 7.223575 | 196.319659 | 0.226804 | 0 | 15.851122 |
| *No_Prev_Dose* | Standard | 103 | 0 | 21.285362 | 164.777883 | 0.145631 | 0 | 0 |
| *No_Prev_Dose* | Aggressive | 97 | 0.597938 | 7.321588 | 200.003069 | 0.391753 | 0 | 17.734339 |
| *No_Receptor_Channel* | Standard | 103 | 0 | 20.827446 | 164.887282 | 0.135922 | 0 | 0 |
| *No_Receptor_Channel* | Aggressive | 97 | 0 | 7.434832 | 277.332284 | 1 | 0.123711 | 11.099976 |

The full model achieved a 74.2% success rate in the aggressive cohort, with mean peak cytokines of 195.7, indicating that it could control tumor burden while keeping toxicity near but below the critical safety region. Removing the class hint caused the largest drop in performance. In the aggressive cohort, success fell from 74.2% to 0%, mean final tumor burden increased from 7.34 to 231.11, and mean peak cytokines rose from 195.7 to 471.4, with universal exceedance of both the 200 and 300 cytokine thresholds. This shows that phenotype information is essential for identifying which patients require active intervention.

Removing the receptor channel also reduced aggressive-cohort success to 0%. Although mean final tumor burden remained low (7.43), mean peak cytokines increased to 277.3, all aggressive patients exceeded the 200 threshold, and some exceeded 300. This indicates that receptor-level information is not only useful for tumor control, but is especially important for keeping toxicity within safe limits. By contrast, removing the delta features had little effect. Performance remained similar to the full model, suggesting that short-term trend information contributed relatively little beyond the current state variables. Removing previous-dose memory caused a moderate decline in performance, reducing aggressive-cohort success to 59.8% and increasing both cumulative dose and cytokine overshoot. This suggests that treatment history helps the controller maintain smoother and safer dosing, but is less critical than phenotype and receptor-state information.

Overall, the ablation results show that the main advantage of OPTIMIS comes from two sources: phenotype-aware control and microscale receptor-state feedback. Together, these features enable the controller to balance tumor suppression and toxicity more effectively than reduced versions of the model.

### Superiority over baselines and heuristic clinical protocols

We compared the final RL_Doctor, which used the full multiscale observation space and learned a closed-loop control policy through PPO policy, against several simpler treatment strategies, as listed in **Table 2**. No_Drug applied zero Dasatinib throughout the full treatment course. Constant_Low_0.2 applied a fixed low dose of 0.2 at every step. Constant_Medium_0.5 applied a fixed medium dose of 0.5 throughout the course. Heuristic_Adaptive used a simple reactive rule-based policy that adjusted dosing according to observed macroscale state variables, primarily tumor burden and cytokine level, without using reinforcement learning or receptor-level feedback.

**Table 2:** Comparative performance of the AI-driven reinforcement learning policy against standard constant and adaptive heuristic dosing strategies in managing tumor burden and cytokine toxicity across distinct patient phenotypes.

| *Policy* | *Cohort* | *Num_Patients* | *Success_Rate* | *Mean_Final_Tumor* | *Mean_Peak_Cytokines* | *Frac_Cytokines_Over_200* |
| --- | --- | --- | --- | --- | --- | --- |
| *Constant_Low_0.2* | Standard | 105 | 0 | 49.483569 | 134.806542 | 0.009524 |
| *Constant_Low_0.2* | Aggressive | 95 | 0 | 497.49281 | 513.018357 | 1 |
| *Constant_Medium_0.5* | Standard | 105 | 0 | 110.650433 | 85.541078 | 0 |
| *Constant_Medium_0.5* | Aggressive | 95 | 0 | 73.489366 | 414.39393 | 1 |
| *No_Drug* | Standard | 105 | 0 | 21.379072 | 167.59782 | 0.095238 |
| *No_Drug* | Aggressive | 95 | 0 | 644.33538 | 544.166931 | 1 |
| *Heuristic_Adaptive* | Standard | 92 | 0 | 24.741293 | 146.505608 | 0.043478 |
| *Heuristic_Adaptive* | Aggressive | 108 | 0 | 549.038037 | 524.88837 | 1 |
| *RL_Doctor* | Standard | 92 | 0 | 21.295475 | 169.638481 | 0.130435 |
| *RL_Doctor* | Aggressive | 108 | 0.712963 | 7.237788 | 200.540295 | 0.287037 |

In the aggressive cohort, all of these simpler strategies performed poorly. The no-drug policy led to severe tumor progression and extreme cytokine toxicity. Fixed-dose strategies also failed to balance efficacy and safety: although they sometimes reduced tumor burden, they consistently drove cytokines far above safe levels and produced 0% success in aggressive patients. The reactive heuristic policy also failed in the aggressive cohort. Because it adjusted treatment only after toxicity became visible at the macroscale, it acted too late to stop the inflammatory cascade. As a result, it showed 0% success and much higher cytokine peaks than the RL policy. This suggests that purely reactive dosing is not sufficient in high-risk disease.

In contrast, the RL_Doctor policy achieved the best overall balance between tumor control and toxicity. In the aggressive cohort, it reduced mean final tumor burden to about 7.3, kept mean peak cytokines near the 200 safety boundary, and achieved a success rate of about 64–74% depending on the evaluation set. It was also the only strategy that consistently avoided the severe cytokine overshoot seen with the simpler baselines. For the standard cohort, the RL policy behaved similarly to the no-drug baseline, using little or no Dasatinib. This indicates that the controller did not simply learn to dose more often; instead, it learned to reserve treatment for cases where intervention was actually needed. Overall, these comparisons show that OPTIMIS outperformed both static schedules and a simple adaptive heuristic, especially in the aggressive cohort where precise, anticipatory control was most important.

## Concluding Discussions

In summary, the development and validation of the OPTIMIS platform demonstrates a fundamental paradigm shift in the computational optimization of cellular therapies. By successfully bridging the stochastic microscale of receptor engagement with macroscopic tissue dynamics, our reinforcement learning agent autonomously revealed that reactive, symptom-driven clinical protocols are mathematically insufficient for high-risk patient phenotypes. The AI’s emergent three-phase “surfing” strategy—characterized by a preemptive pharmacological brake, a controlled taper, and a precise soft-landing pulse—highlights a critical biological insight: systemic cytokine elevation is a lagging indicator of a cellular cascade that has already achieved dangerous exponential momentum. Our ablation studies indicate that microscale receptor activity functions as an important early-warning computational biomarker within the OPTIMIS framework, particularly for preventing toxicity overshoot in aggressive disease.

The computational novelty lies in the integration of three distinct components into a single multiscale control architecture. First, the platform preserves mechanistic detail at the receptor/synapse level. This gives the framework a biologically interpretable microscale module rather than treating all dynamics as a purely black-box process. Second, the platform learns a drug-aware neural ODE surrogate for macro-scale tumor, CAR-T, and cytokine dynamics from synthetic multiscale trajectories. This creates a differentiable and computationally efficient digital twin that captures the system-level evolution of disease and treatment response while remaining coupled to mechanistic receptor-state updates. Third, the platform embeds this multiscale digital twin inside a reinforcement learning environment, allowing a controller to learn adaptive dosing policies that explicitly trade off tumor control against toxicity and treatment burden. The RL agent does not optimize a static regimen; it learns a sequential decision policy over time. Taken together, the platform is novel because it combines mechanistic microscale simulation, learned macro-scale surrogate dynamics, and adaptive reinforcement learning control in a single persistent framework for therapy design.

The most immediate impact of this platform is as an in silico testbed for adaptive CAR-T treatment design. It can be used to compare dynamic dosing strategies against simpler schedules such as no drug, constant drug, or rule-based therapy. This supports hypothesis generation about when and how signaling-modulating drugs should be used during CAR-T treatment. A second potential impact is in patient stratification and personalized control. Because the framework can encode phenotype-dependent dynamics, it can be used to explore whether aggressive and standard disease states require different dosing strategies, toxicity thresholds, or receptor-control regimes. In future extensions, the same logic could be applied to more realistic patient-specific parameterizations. A third impact is in mechanistic interpretability. Since the platform explicitly tracks receptor activation, tumor burden, CAR-T abundance, cytokines, and drug dosing, it can help identify why a policy works, not just whether it works. This is valuable for translational research, where mechanistic plausibility matters in addition to predictive performance. A fourth impact is in preclinical optimization and simulation-guided design. Before moving to animal models or clinical trials, investigators could use such a platform to screen candidate intervention rules, stress-test toxicity-control strategies, and identify regimes that appear robust across simulated patient heterogeneity. More broadly, the framework could be generalized beyond CAR-T therapy. The same computational strategy could be applied to other biomedical settings where multiscale dynamics and adaptive control are central, including immune modulation, targeted cancer therapy, infectious disease treatment, and combination therapy scheduling.

## Code availability

All relevant source codes can be found in the GitHub repository: https://github.com/wujah/OPTIMIS.

## Acknowledgement

This work was supported by the National Institutes of Health under Grant Numbers R01GM122804, and the United States – Israel Binational Science Foundation Project Number: 2023336. The work is also partially supported by a start-up grant from Albert Einstein College of Medicine. Computational support was provided by Albert Einstein College of Medicine High Performance Computing Center. Finally, we also greatly appreciate the general support from the Einstein 2030 Seed Fund.

## Author Contributions

Z.S. and Y.W. designed research; Z.S. and Y.W. performed research; Z.S., and Y.W. analyzed data; Z.S. and Y.W. wrote the paper.

